# Motile bacteria collectively transport soil water during host colonisation

**DOI:** 10.64898/2026.09.02.748768

**Authors:** IC Engelhardt, J Anguita, D Barriales, TJ Daniell, N Holden, LX Dupuy

## Abstract

Nutrient availability in soil is temporally and spatially heterogeneous, and, as a result, microbial migration is critical for many species. The nature of microbial movement in soil, however, is unknown due to a lack of observations and experimental data. We developed live-imaging and image-analysis techniques to track the movement of single cells through soil to elucidate how *Bacillus subtilis* utilises pore space during the early root colonisation. The study reveals that the bacterium can modify fluid pathways to create streams, even at low bulk cell density. The phenomenon was influenced by pore structure, distance from the root and the viscosity of the soil solution. By generating macroscopic fluid motion, bacteria may also be able to travel faster and farther than individually, while limiting energy expenditure.

## Main Text

Bacteria are known to possess sophisticated mechanisms to locate and navigate toward nutrient sources within the environment. These include a broad diversity of chemoreceptors, chemotaxis, haptotaxis, various forms of active motility, and complex systems for cooperation between cells (quorum sensing).

In soil, sensing of and movement towards nutrient sources is particularly challenging for microorganisms since it represents a complex matrix of physical obstacles, heterogeneous nutrient distribution, and variable pore-space connectivity. The impact of texture and hydration status on bacterial mobility (Fontes *et al.,* 1991; Ebrahimi and Or, 2014; Tecon and Or, 2016) and diversity (Zhou et al., 2002; Nunan *et al.,* 2004) is frequently highlighted. Diffusion of chemoattractant through soil is heterogeneous, and the presence of solid surfaces can significantly alter the chemotactic behaviour of bacteria (Yang *et al.,* 2019). Above all, soil is a relatively carbon-poor environment in which inefficient energy expenditure can dramatically affect survival (Hobbie and Hobbie, 2013). Live observations of soil microbes are particularly scarce, and knowledge of how bacteria utilise different motility strategies to navigate the soil matrix host is thus limited.

Here we used custom-made live imaging systems and image analysis to track the movement of single bacterial cells in the soil pore network. Transparent soil microcosms (see Lui et al., 2021 for details) were saturated in Hoagland’s solution (C-free plant nutrient solution) diluted in either a high-viscosity colloid (Percoll, GE Healthcare) or water (low viscosity). To track fluid movement in the pore space, the microcosms were inoculated with green fluorescent microspheres (Sigma-Aldrich) and/or approximately 2 million *B. subtilis* cells (red fluorescent NRS5852 strain; see supplementary materials for details and preparation).

Imaging was performed on a Leica SP08 TCS confocal microscope, and particle Image Velocimetry (PIV) using the iterative Lucas-Kanade solver (Le Besnerais and Champagnat 2005) was used to track the movement of bacteria and fluorescent microspheres in image stacks. The root was manually selected and traced from the auto-fluorescence signal, while Weka segmentation algorithms were used to automatically identify soil particles and pore space (Fig. S1A-C). Distance to root and soil particles was determined using geometry-to-distance maps (Fig. S1D-E). Pore size was determined as the largest sphere fitting in the space using the local thickness function (Fig. S1F).

RStudio (R Core Team 3.1.2) was used to run all ANOVA analyses of significance, post-hoc tests (Tukey), correlation plots and linear regressions on bacterial and microsphere data. Segmented quantile regression (Hao *et al*., 2017; Mills *et al*., 2006; Mills *et al*., 2009) evaluated the impact of environmental factors on bacterial density estimated from fluorescence intensity (Fig. S2) and velocity estimated with PIV.

In dual-inoculation experiments (motile *B.subtilis* with static microbeads), we observed cell acceleration at bottlenecks and slowing as the pore space enlarged (Fig. 1A). The velocity profile in the pore space was parabolic (Fig. 1B), consistent with the Hagen– Poiseuille law for laminar flow through cylindrical pipes (Kirby 2010). We confirmed that such collective movements are linked to streams of water in the pores by correlating the magnitude and direction of the cell velocity with that of fluorescent microspheres (Fig. S3). The average velocity profiles of the bacterial and microsphere flows were comparable, with the maximum velocity near the centre of the flow, and the average velocity of bacteria usually exceeding that of microspheres. We found that the presence of motile bacteria was the most important factor (more than root water uptake or other convection) driving fluid movement in the pore space in our system (Fig. 1C; Table S1).

**Fig 1:**
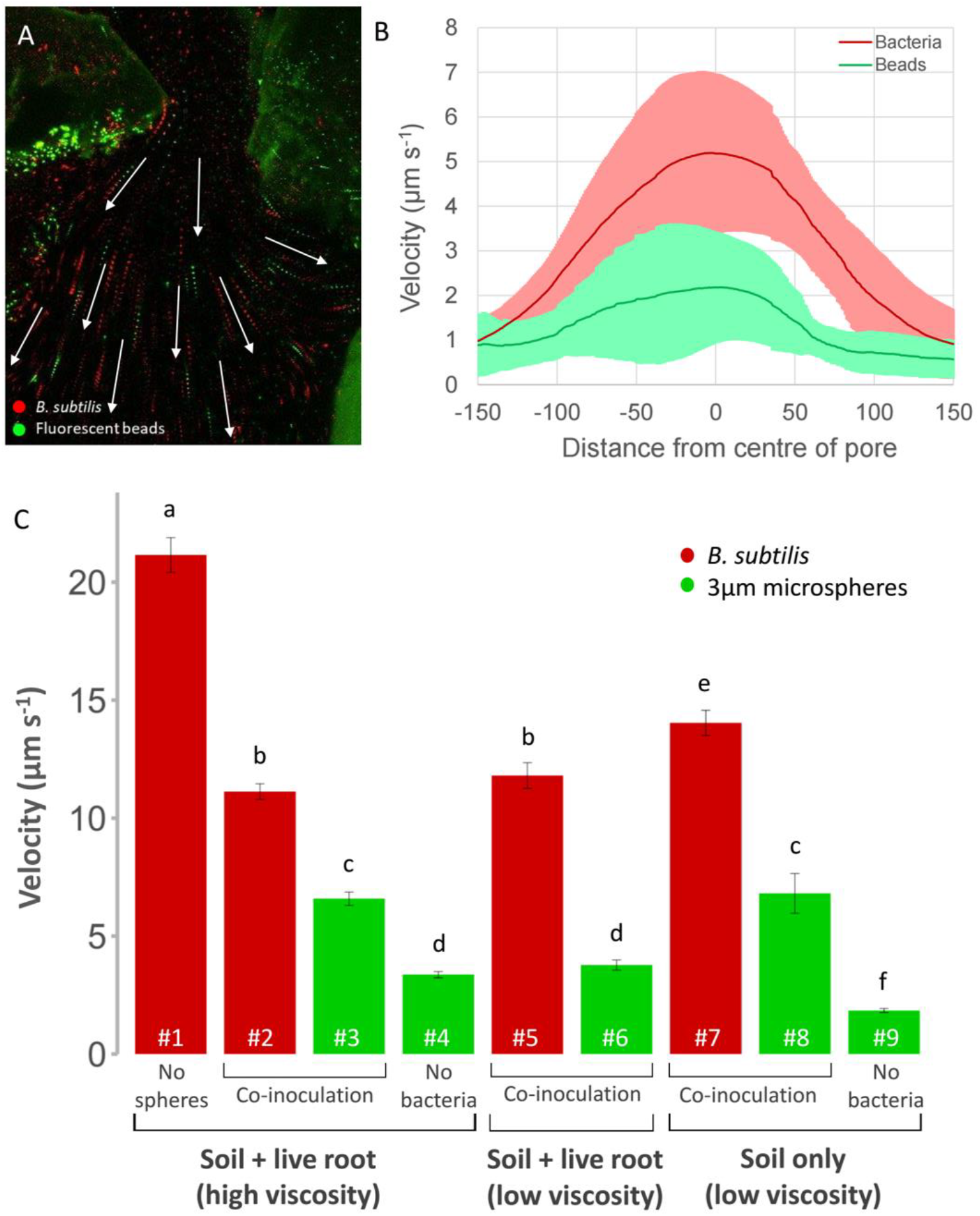
Bacterial movement through the pore space cause active fluids within the soil. (A) The trajectories of immobile, fluorescent microspheres (3um diameter, green) align with the bacterial trajectories (*B. subtilis*, red) in the pore space around growing wheat roots. (B) The velocity profile of beads (green) mimic the velocity profile of bacteria (red), at perpendicular transects to flow direction though several pores. Solid lines show the average velocity profile for each bacteria and microspheres with the lighter shade polygon representing ±SD. The velocity and angle of bead trajectories in the soil are compared to their closest bacterial neighbour. The correlation between bead and bacterial velocities (C-D) as well as the angle of the bead and bacterial trajectories (E-F) were examined under 2 levels of viscosity, high (C and E) and low (D and E). The r2values indicate the f fit of the linear correlation and *** indicates a significance level at P < 0.05.

Viscosity can affect bacterial movement by increasing drag forces on swimming bacteria and indirectly influencing the diffusion of molecules such as nutrients or surfactants. Generally, bacterial swimming velocity increases with increasing viscosity up to a threshold (approximately 2 mPa s^-1^) after which increasing viscosity becomes a limiting factor (Kaiser and Droetsch 1975; Schneider and Droetsch 1974; Shoesmith 1960). The high-viscosity (6.6 mPa s-1) colloid used in this experiment is well within the range associated with movement impediments, but moving speeds were not significantly reduced. The positive correlation between bacterial and microsphere velocity was unaffected by the increase in viscosity (Fig. S3), but the average velocity of the co-inoculated microspheres was reduced (Fig. 1), while their alignment to bacterial cells increased (Figure S3 C-D) with r^2^ = 0.52 (p < 0.05) and r^2^ = 0.94 (*p* < 0.05) for low and high viscosity, respectively. Movement of water resulting from the motility of soil microbes, termed active fluids here, could be a critical mechanism in natural soils where fluctuations in viscosity, brought on by the rapid changes in water saturation (Lee *et al.,* 2013) may occur. This phenomenon could represent an energy-efficient mechanism for moving through mechanically limiting environments, even at lower cell densities.

The non-monotonic “hump-shape” curve resulting from the best-fit 3rd-order polynomial (Fig. 2A, r^2^ = 0.53, p < 0.05) through the boundary points in the bacterial velocity vs pore size scatter suggests a limit (about 500 µm diameter) beyond which formation of active fluids in soil is not possible, which confirms previous finding that collective movement of motile bacteria is strongly impacted by confinement (Lushi *et al.,* 2014), and spatio-temporal organisation is driven by boundaries and by bacterial interactions with them (Wioland *et al.,* 2013). Our results suggest a potential optimal pore size for low-density bacterial migration, which may lie at the trade-off between movement-induced shear stress and the requirement for a critical degree of bacterial alignment to initiate movement.

**Fig 2:**
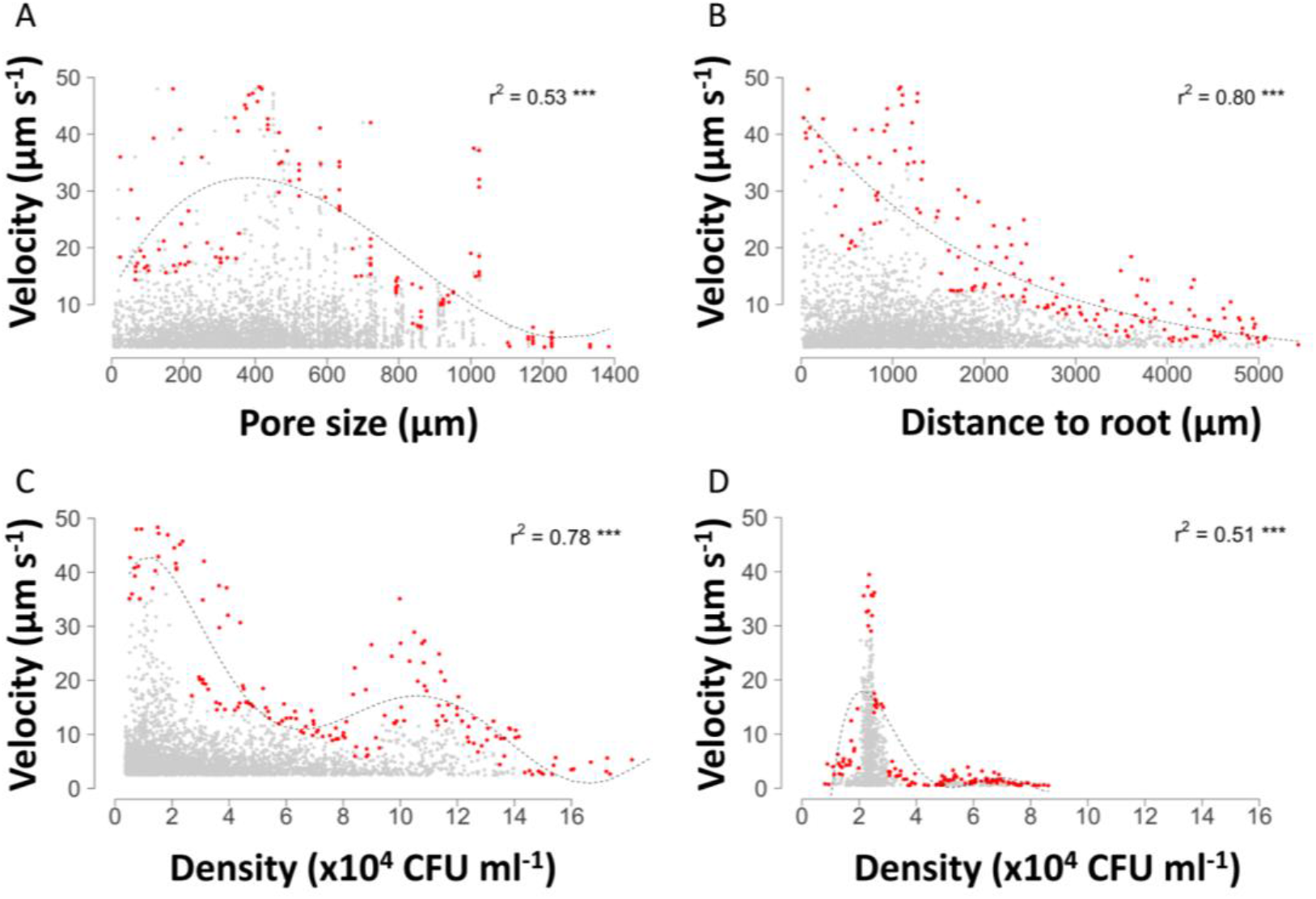
Environmental factors such as soil physical attributes, presence of live roots and bacterial cell density govern bacterial flows. Scatter plots (grey), boundary points (red) and constraint lines (black) show the impact of pore size (A) and distance to root (B) on bacterial velocity. The unit for bacterial velocity is in µm. s-1while pore size and distance to root are both in µm. Boundary points of the bacterial velocity vs density scatter show 2 peaks in bacterial velocity at different cell density ranges when in the presence of a live plant root (C) and a single peak in bacterial velocity when in the absence of live plants (D). The regression models of the boundary points were chosen from straight linear lines or 2ndand 3rdorder polynomials or exponential curves based on best fit (r2value) and *** indicates significance of the fit at p < 0.05.

The presence of the root also appeared to stimulate the formation of active fluids, with bacteria closest to the root reaching the highest velocities (Fig. 2B, r^2^ = 0.80, p < 0.05). Additionally, two peaks in the bacterial velocity were identified in the presence of live roots (Fig. 2C, r^2 =^ 0.73, *p* < 0.05). A group of bacteria acquired high velocities (max 40 - 50 µm s-1) at low densities (∼2 × 10^4^ cells ml-1), and a group acquired high velocities (max 30 - 40 µm s^-1^) at higher densities (∼1 × 10^5^ cells ml^-1^), which highlights the likely dependency of motility on cell density. Both peaks in bacterial velocities were found to be significantly closer to the root (< 2mm) than the low-velocity groups (Fig. S5). Since only one peak at low cell density was detected when the root is replaced with nutrient solution (Fig. 2D, r^2 =^ 0.51, *p* < 0.05), we deduce that root dynamics drive the high velocity at high cell density in our system. When colonising a host, bacteria must utilise perceived nutrient gradients to orient towards nutrient sources, which appears to result in the directed flow of active fluids. The peak in fluid velocities observed at higher bacterial density in the presence of plant roots may be due to chemotactically directed active fluids within an otherwise nutritionally heterogeneous environment.

At very high cell densities such as in bacterial swarms or concentrated bacterial suspensions, the movement of self-propelled individual cells results in large-scale coherence or collective fluid motion. Several studies have described a critical cell density at which active fluids occur in concentrated bacterial suspensions (Subramanian and Koch 2009; Ryan *et al.,* 2011). The concentrations in these studies are at least about 10^9^ cells per mL (Wolgemuth 2008; Ariel *et al.,* 2018, Solokov *et al.,* 2007). Here, we present evidence of active fluids driven by bacteria in the soil pore space at cell densities as low as 10^4^ cells mL^-1^ and recorded maximum velocities of up to 50 µm s^-1^ under challenging environmental conditions. It is generally accepted that bacterial individual speeds fall in the region of 15 - 20 µm s^-1^ (Hambey *et al.,* 2018; Voutc‘h *et al.,* 2020), and under controlled laboratory conditions with an abundance of easily available nutrients and at optimum temperatures, maximum speed of almost 25 µm s^-1^ have been recorded in the “run” phase for *B. subtilis* (Turner *et al.,* 2016; Ito *et al.,* 2005; Hambey *et al.,* 2018). In concentrated bacterial suspensions, collective flow speeds have been shown to reach higher velocities, up to 50-100 µm s^-1^ (Sokolov et al. 2007). Collective swimming reaches speeds far higher than that of the individual (Solokov *et al.,* 2007; Ariel *et al.,* 2018; Cisneros *et al.,* 2007; Wolgemuth 2008), which indicates it may be critical for movement in complex environments such as soil where nutrient limitation or physical constraints may be present.

We conclude that, by creating active fluids within soil pores, bacteria may be able to migrate faster and further through challenging environments through a combination of energy-saving and protective. Although affected by pore size and nutrient availability, active bacterial migration through soil may be more extensive than previously estimated, potentially significantly altering our understanding of soil ecosystems. However, in natural conditions, the distance travelled by collective motion may still be severely constrained by nutrient availability, limits in the diffusion of signalling molecules, and reduced connectivity due to the presence of air pockets, with a subsequent requirement to move through water films at the surface of particles.

## Data availability

All data supporting the findings of this study are available within the article and its figures

## Research funding

This work was funded by the European Research Council (ERC) under the European Union’s Horizon 2020 research and innovation programme (Grant agreement No. 647857-SENSOILS). We also acknowledge the funding from the Spanish Ministry of Science and Innovation (MICINN) under de project MICROCROWD (PID2020-112950RR-I00).

## Author contribution

Funding Acquisition: LD. Conceptualisation: LXD, TJD, NH, IE. Experimental Work: IE, BD, AJ. Data Analysis: IE, LD. Supervision: NH, TJD, LXD. Writing – original draft: IE, LXD. Writing – review & editing: NH, TJD, LXD.

