## Supplementary Information for "Motile bacteria collectively transport soil water during host colonisation"

*Live imaging system*

Microcosms consisting of 2 microscope slides separated by a 3-sided PDMS spacer (internal dimensions of 50 x 20 x 4 mm) were filled with sterile Nafion (Lui et al., 2021 for details). Seedlings with a root length of 2-5 mm were transplanted into the Nafion-filled microcosms. Microcosms were saturated in Hoagland's solution (C-free plant nutrient solution) diluted in either a Percoll solution (GE Healthcare), a biologically non-toxic colloid solution (high viscosity), or water (low viscosity). To track the movement of fluids in the pore space, the microcosms were inoculated with fluorescent microspheres (Micro Particles based on melamine resin, FITC marked, Sigma-Aldrich).

*Strain details and preparation*

All *B. subtilis* strains used in this study were derivatives of isolate NCIB 3610 and include NRS1473 (NCIB 3610 sacA::Phy-spank-gfp mut2 (kan) (Verhamme et al, 2007)) and NRS5852 (NCIB 3610 amyE::Phy-spank-mKate (spc), made by Margarita Kalamara, NSW laboratory using a strain from Van Gestel *et al*, 2014. Strains were grown in MSgg medium (5 mM potassium phosphate and 100 mM MOPs adjusted to pH 7.0 and supplemented with 2 mM MgCl2, 700 µMCaCl2, 50 µM MnCl2, 50 µM FeCl3, 1 µM ZnCl2, 2µM thiamine, 0.5% (v/v) glycerol, and 0.5% (w/v) glutamate) for about 24 hours at 20°C, whilst shaking. After incubation, the MSgg solution was replaced with nutrient-free Percoll (or carbon-free Hoeglands solution in viscosity studies) by centrifugation and reconstitution of the bacterial pellet.

*Plant*

Wheat seeds (*Triticum aestivum* var. filum) were sterilised by soaking for 15 minutes in 2% Calcium hypochloride solution followed by thorough rinsing using sterile distilled water and germinating for 1-2 days on 0.5% distilled water agar at 20°C.

*Environmental envelope details*

The data of the environmental factor was binned into 17 to 20 bins. From each bin, the maximum bacterial response variable values were picked (n=10 per bin) to define the outer data envelope boundary points. Several regression models were run on the boundary points (straight line, exponential curve, 2$^{nd}$ order polynomial curve). The best fit model for each environmental factor was chosen based on the $r^2$ and p-value output. Non-linear models were also run using a self-starting function (sslogis) in R.

*References*

**Liu Y,** Patko D, Engelhardt IC, George TS, Stanley-Wall NP, Ladmiral V, Ameduri B, Daniell TJ, Holden N, MacDonald MP, Dupuy LX. Whole plant-environment microscopy reveals how *Bacillus subtilis* utilises the soil pore space to colonise plant roots. *Proceedings of the National Academy of Sciences of the United States of America*. 2021; 118(48).

**Verhamme DT**, Kiley TB, Stanley-Wall NR. DegU Co-Ordinates Multicellular Behaviour Exhibited by *Bacillus Subtilis*. *Molecular Microbiology*, 2007; 65 (2): 554–68.

**van Gestel J**, Weissing FJ, Kuipers OP, Kovács ÁT. Density of founder cells affects spatial pattern formation and cooperation in Bacillus subtilis biofilms. *ISME Journal*, 2014; *8*(10), 2069–2079.

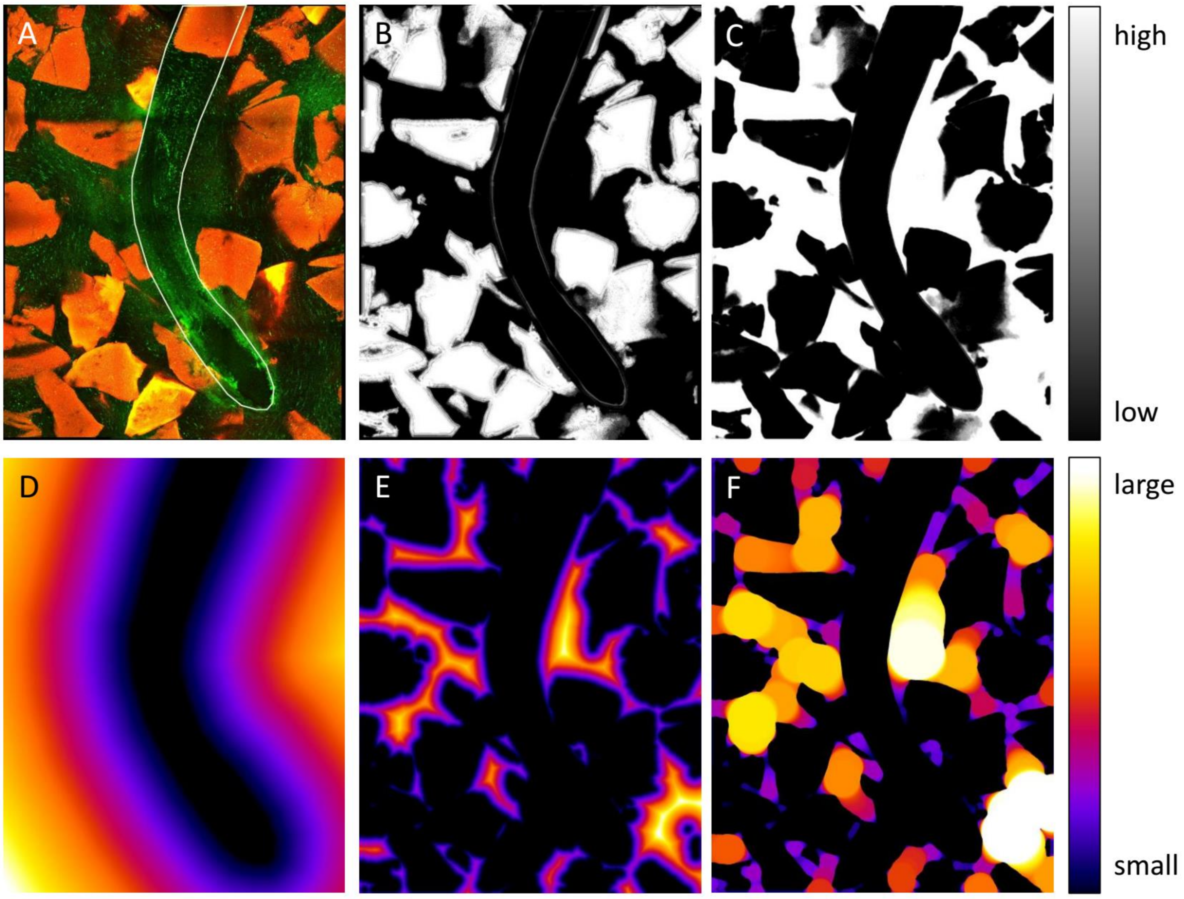


**Fig S1: Image analysis pipeline.** (A) The intensity of the GFP signal was increased so that the root could be manually traced using the natural auto-fluorescent signal. The nafion particles are shown in red and the bacteria and root are seen in green. Weka segmentation in image j was used to automatically predict probability of (B) nafion particles and (C) pore space. Probability maps are represented in grey scale from high probability (white) to low probability (black) of classifier. Geometric distance maps were created in image j using the (D) manual root tracing and (E) pore space probability map. (F) Local thickness function in image J was used to determine the pore size throughout the binary probability image. All distance maps were used a colour scale from blue (small distance/ sphere) to white (large distance/ sphere).

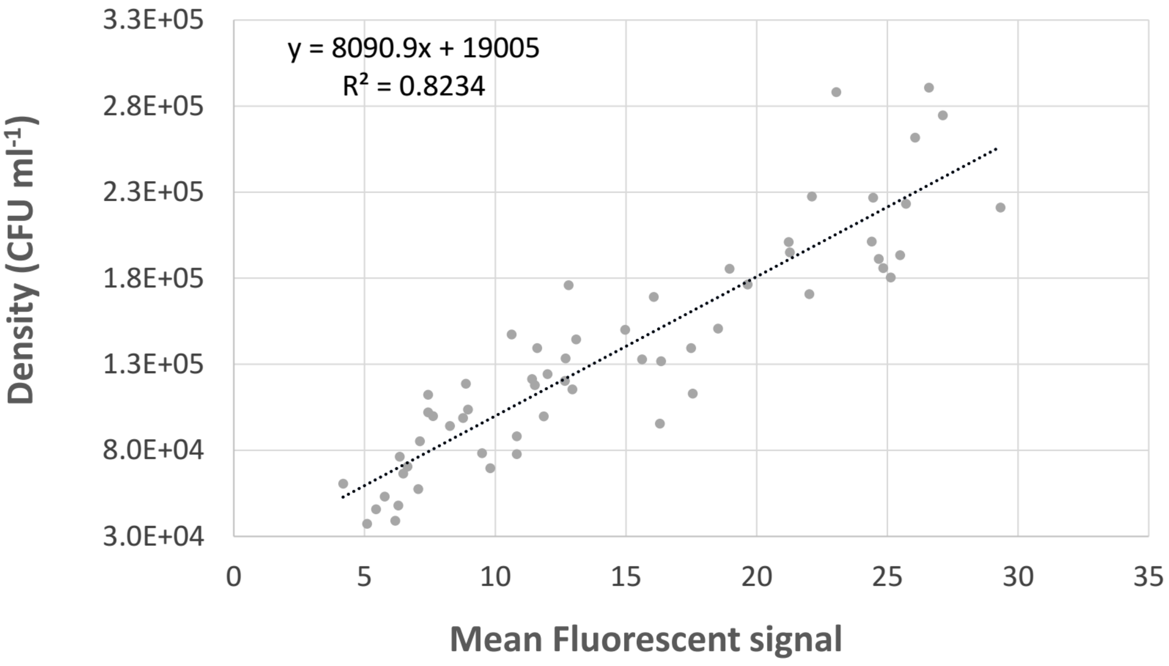


**Fig S2: System calibration for cell density based on mean fluorescent signal.** A total of n=59 regions where selected on 19 separate composite volume images obtained from at least 9 individual live systems at different time points. The number of trajectories were counted, the average fluorescent signal measures and the volume of calculated for each of the selected regions. The bacterial density as trajectories per volume of pore space (CFU per ml) are plotted against the mean fluorescent signal in the region. The linear correlation was used to calibrate the bacterial density measures. The peak bacterial densities captured in the active fluids lie around 1.5 x $10^5$ thus well within the range included in this calibration curve.

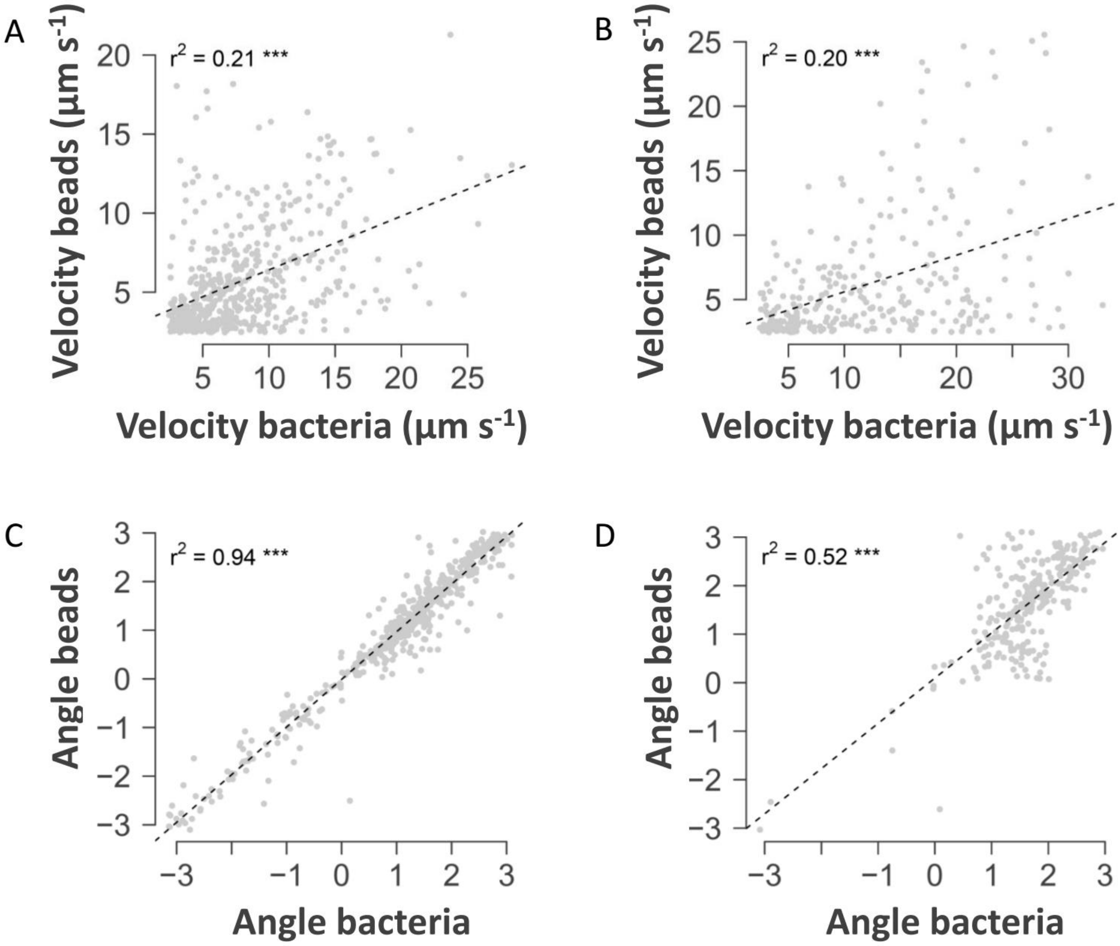


**Fig S3: The velocity and angle of bead trajectories in the soil compared to that of their closest bacterial neighbour**. The correlation between bead and bacterial velocities (A-B) as well as the angle of the bead and bacterial trajectories (C-D) were examined under 2 levels of viscosity, high (A and C) and low (B and D). The r 2 values indicate the f fit of the linear correlation and *** indicates a significance level at P < 0.05.

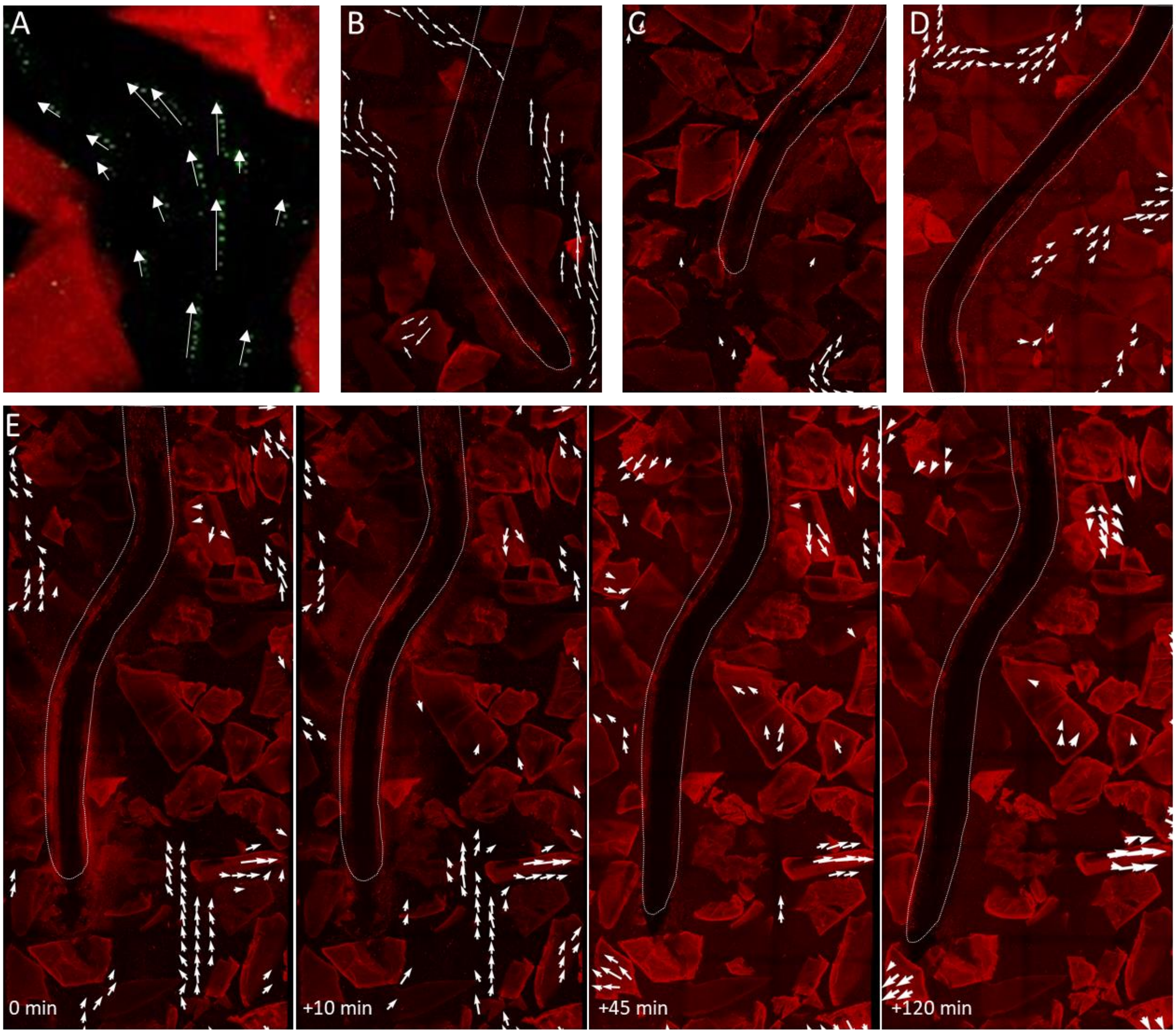


**Fig S4: Bacterial trajectories in the soil pores around live plant roots**. (A) *B. subtilis* cells captured in image stacks form trajectories in the pore space. (B-E) PIV detects particle movement and assigns vectors with direction (arrowhead) and magnitude (arrow size). Bacterial flows through the pore space can be (B) long and well connected or (C) short and disconnected. (D) Bacteria navigate the pore space with flows merging or splitting around obstacles. (E) Image sequence of the same root show bacterial flows in the soil space are dynamic, changing paths in short time spans.

| Finding | Bars | significance |
|---|---|---|
| Movement of static microbeads increased in the presence of plant roots. | #9 vs #4 | $p < 0.05$ |
| Movement of static microbeads increased in the presence of motile bacteria. | #8 vs #9 | $p < 0.05$ |
| Bacteria increase microbead movement significantly more than plant roots | #4 vs #8 | $p < 0.05$ |
| When motile bacteria are present, there is no significant effect of plant roots on microbead movement. | #3 vs #8 | $p = 0.999$ |
| The velocity of active fluids around plant roots is not affected by the tested viscosity levels. | #2 vs #5 | $p = 0.577$ |
| The velocity of microspheres (in the presence of bacteria) is reduced at low viscosity. | #3 vs #6 | $p < 0.05$ |
| The velocity of bacteria is higher in a defined bacterial solution than with sole reliance on root exudate. | #2 vs #7 | $p < 0.05$ |
| The velocity of microspheres is not affected by the nutrient source. | #3 vs #8 | p = 0.999 |

**Table S1: D**etails of the pairwise ANOVA highlighting the individual findings, reference to pairs compared based on the bar graph (Fig. 1C) and significance of each.

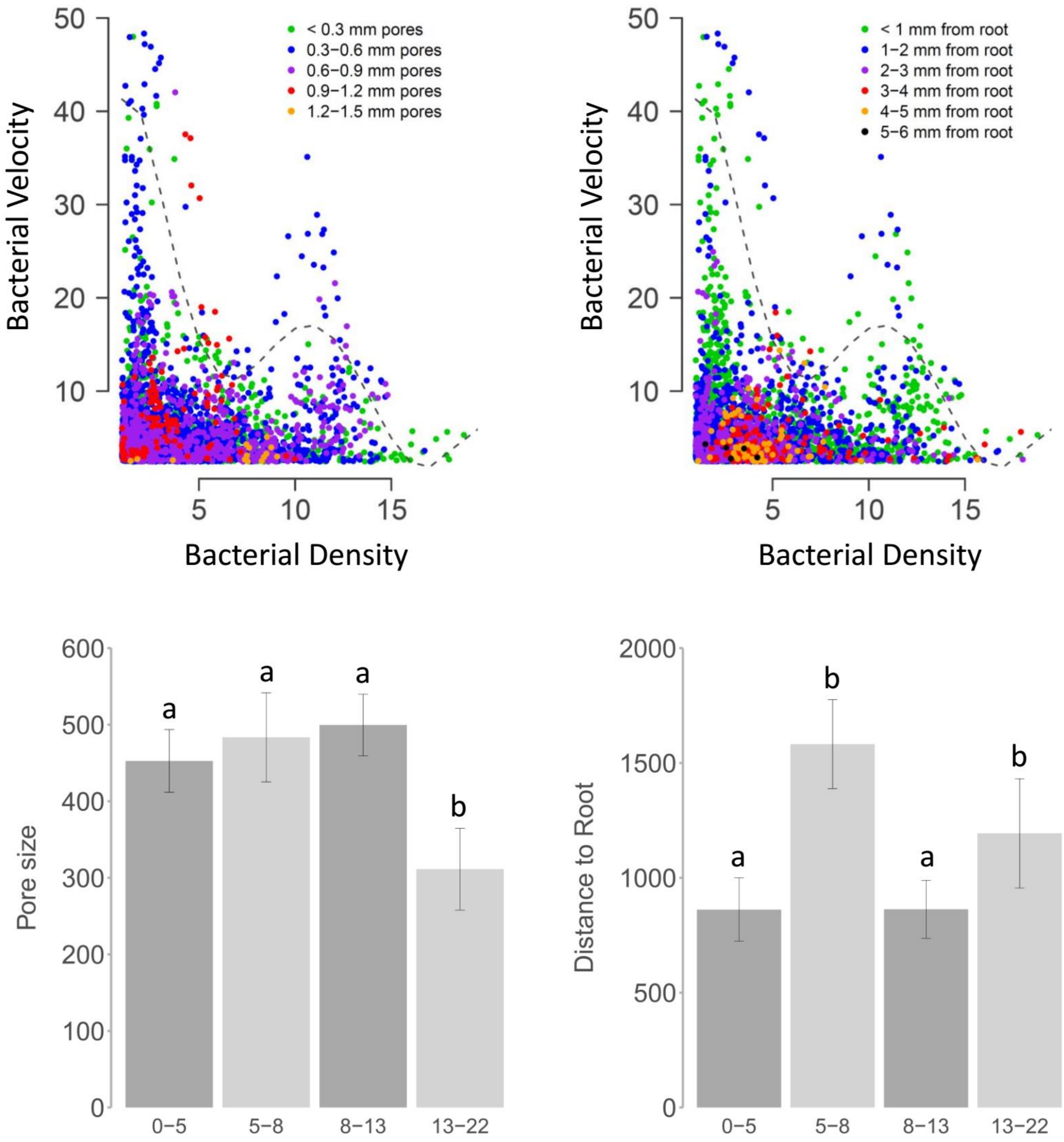


**Fig S5: The correlation between bacterial density and bacterial velocity around live roots is governed by several factors.** Boundary points (red) of the bacterial velocity vs density scatter (grey) have a best fit 5th order polynomial regression (top right, $p>0.05$, $r2=0.73$) with 2 peaks in bacterial velocity at different densities (top left). 4 different groups of bacterial response were identified within the boundary points along the density gradient (top right); a high velocity group at low bacterial density (purple), a low velocity group at low density (green), a high velocity group at high density (red) and a low velocity group at high density (blue). The pore size and root proximity distribution within the 4 bacterial groups are shown in the bar plot. For each graph separately, groups which share a letter are not significantly different from each other. Significance is at $P < 0.05$ level. Error bars show the 95% confidence interval.
